# Persistent Chikungunya Virus Replication in Human Cells is Associated with Presence of Stable Cytoplasmic Granules Containing Non-structural Protein 3

**DOI:** 10.1101/236703

**Authors:** Roland Remenyi, Yanni Gao, Ruth E Hughes, Alistair Curd, Carsten Zothner, Michelle Peckham, Andres Merits, Mark Harris

## Abstract

Chikungunya virus (CHIKV), a mosquito-borne human pathogen, causes a disabling disease characterized by severe joint pain that can persist for weeks, months or even years in patients. The non-structural protein 3 (nsP3) plays essential roles during acute infection, but little is known about the function of nsP3 during chronic disease. Here, we used sub-diffraction multi-color microscopy for a spatial and temporal analysis of CHIKV nsP3 within human cells that persistently replicate viral RNA. Round cytoplasmic granules of various sizes (i) contained nsP3 and G3BP Stress Granule Assembly factor; (ii) were next to double-stranded RNA foci, and nsP1-positive structures; and (iii) made contact with markers of the cytoskeleton and cellular structures, such as early endosomes and nucleopores. Analysis of protein turnover and mobility by live-cell microscopy revealed that granules could persist for hours to days, can accumulate newly synthesized protein, and move at differently through the cytoplasm. Granules also had a static internal architecture and were stable in cell lysates. Whereas cells with active replication and stable nsP3-granules did not respond to oxidative stress, refractory cells that had cleared the non-cytotoxic replicon could. In summary, nsP3 can form uniquely stable granular structures that persist long-term within the host cell. This continued presence of viral and cellular protein-complexes has implications for the study of the pathogenic consequences of lingering CHIKV infection and the development of strategies to mitigate the burden of chronic musculoskeletal disease brought about by a medically important arthropod-borne virus (arbovirus).

**Importance:** Chikungunya virus (CHIKV) is a re-emerging alphavirus transmitted by mosquitos and causes widespread transient sickness but also chronic disease affecting muscles and joints. Although no approved vaccines or antivirals are available, a better understanding of the viral life cycle and the role of individual viral proteins can aid in identifying new therapeutic targets. Advances in microscopy and persistent CHIKV model systems now allow researchers to study viral proteins within controlled laboratory environments. Here we established human cells that stably replicate viral RNA and express a tagged version of non-structural protein 3. The ability to track this viral protein within the host cell and during persistent replication can benefit fundamental research efforts to better understand long-term consequences of the persistence of viral protein complexes and thereby provide the foundation for new therapeutic targets to control CHIKV infection and treat chronic disease symptoms.

## Introduction

Chikungunya virus (CHIKV), a re-emerging arbovirus of the Alphavirus genus, causes a transient illness with debilitating symptoms (fever, headache, rash, myalgia, and arthralgia). Chronic infection is common and joint pain can persist for months to years (1–3). Half of the patients from the recent Latin American outbreak may develop chronic inflammatory rheumatism, raising the health burden of musculoskeletal disease in endemic areas (4, 5). During acute infection, CHIKV induces cytopathic effects and apoptosis leading to direct tissue injury and local inflammation (6–8). Biopsies have also revealed the persistence of CHIKV antigens and RNA in synovial macrophages and muscle tissue (1, 9). CHIKV also persists in mice and non-human primate models (10–13). Chronic disease may be a consequence of persistent, replicating and transcriptionally active CHIKV RNA (13), but an understanding of CHIKV’s long-term effects is still emerging.

The ~12-kb positive-sense RNA genome of CHIKV encodes four non-structural proteins, nsP1 to nsP4, which make up the viral replication and transcription complex (reviewed in (14)). A subgenomic RNA expresses six structural proteins. Cellular responses to infection include apoptosis, interferon signalling, stress granule (SG) formation, unfolded protein response, host cell shut-off, and autophagy (reviewed in (15)). Previous research on alphaviruses established the vital role that nsP3 plays in counteracting cellular responses (16–20) and identified essential protein-protein interactions between nsP3 and host proteins (16, 21–23). However, few studies have systematically investigated the long-term effects of persistently replicating CHIKV RNA and continued expression of proteins such as nsP3 on human cells. Although recent studies characterize the formation of organelles that contain nsP3 during acute infection and transient replication (16, 24–27), a corresponding characterization during persistent CHIKV replication is missing. To address these gaps, we sought to further develop CHIKV replicons capable of persistent replication in human cells and to harness this system for analysis by sub-diffraction multi-color microscopy.

We previously characterized transient viral replication in mammalian and invertebrate cell lines (27) and tagged nsP3 with the versatile SNAP-tag for advanced fluorescence microscopy applications (26). The development of a non-cytotoxic CHIKV replicon allowed the establishment of persistent replication in a human cell line (28). Here, we extended the SNAP-based labeling system to this non-cytotoxic CHIKV replicon and generated a human cell line that persistently replicates viral RNA and stably expresses SNAP-tagged nsP3. We then characterized nsP3-containing cytoplasmic granular organelles by sub-diffraction multi-color microscopy. The nsP3-containing granules overlapped with G3BP and were near double-stranded RNA (dsRNA) foci and nsP1-positive structures; moreover, they appeared associated with the cytoskeleton and cellular structures such as early endosomes and nucleopores. Granules persisted for hours and days, can accumulate newly synthesized protein, and move with different speeds through the cytoplasm. Granules did not dynamically exchange SNAP-nsP3 with their surroundings and were stable in cell lysates. Whereas cells with active replication and stable nsP3-granules did not respond to oxidative stress, cells that had cleared the non-cytotoxic replicon could form SGs. In summary, this study aims to contribute to a growing area of research on virus-host interactions during CHIKV infection by coupling a sub-diffraction microscopy analysis with an improved system to track nsP3 during persistent replication. This report is the first to shed light on the persistence of stable intracellular granules of nsP3 within human cells. In turn, understanding the link between the persistence of stable viral protein complexes and pathogenesis has relevance to future studies of chronic CHIKV disease.

## Results

### Development of a stable human-origin cell line carrying a SNAP-tagged CHIKV replicon and super-resolution microscopy of nsP3-G3BP-granules

To determine the intracellular distribution of nsP3, we previously generated a SNAP-tagged replicon (26). Whereas this replicon is cytotoxic and replicates transiently, non-cytotoxic replicons can establish persistent replication in the human cell line HuH-7 (28). To improve the HuH-7 CHIKV cell line, we added a SNAP-tagged nsP3 to a non-cytotoxic replicon and selected puromycin-resistant cells, which will be called “stable CHIKV cells” throughout this paper. Silicon-rhodamine-conjugated O6-benzylguanine probes (BG-647 SiR) labeled SNAP-nsP3 and revealed nsP3-granules (Fig. 1A) comparable to those formed by CHIKV^P3-SNAP^ and CHIKV^P3-ZsGreen^, viruses harbouring SNAP- or ZsGreen-tagged nsP3 (Fig.1B-C). Further experiments focused on the characterization of these nsP3-granules. Whereas cells infected with CHIKV^P3-ZsGreen^ only displayed a granular nsP3-ZsGreen distribution pattern, cells infected with CHIKV^P3-SNAP^ also made rod-like structures (Fig. S2A) as was described previously (26, 27). However, the presence of rods did not correlate with infectivity, as ZsGreen-and SNAP-tagged viruses replicated to similar titres (Table S1).

**Figure 1.**
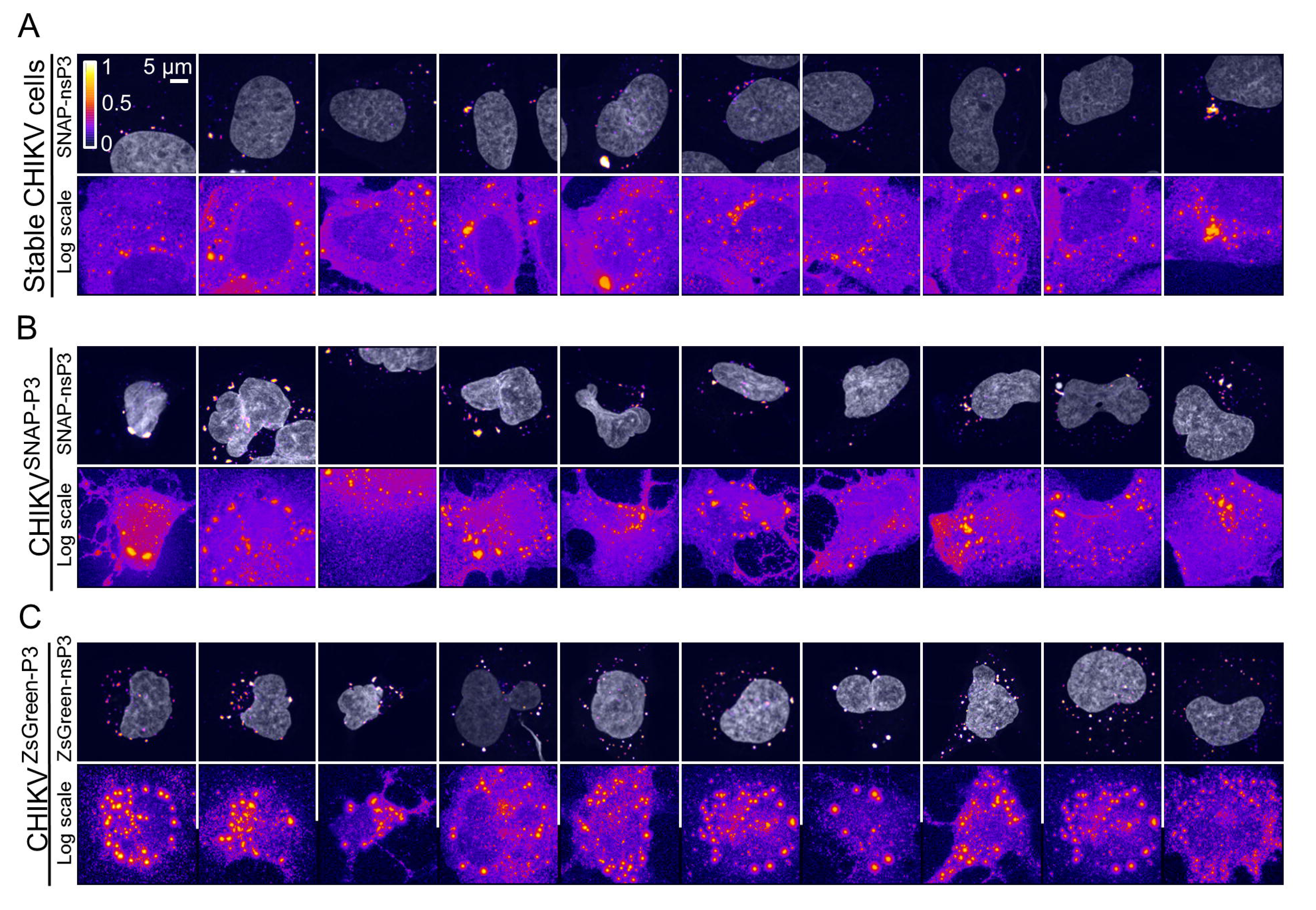
NsP3 has a granular distribution in stable CHIKV cells and infected HuH-7 cells. Top panels: Sub-diffraction confocal microscopy of BG-647-SiR-labeled, imaged in far-red channel. Stable CHIKV cells were chemically fixed and stained with fluorescent BG-647 SiR, which irreversibly binds SNAP-tagged proteins. For CHIKV infection studies, naïve HuH-7 were infected with viral stocks of CHIKV^SNAP-P3^ or CHIKV^ZsGreen-P3^ at an MOI of 10. For cells infected CHIKV^ZsGreen-P3^, no additional labelling was necessary and images were acquired in the green channel. Data shown are maximum-intensity-projections of Z-stacks acquired on an Airyscan confocal system, operated in the super-resolution mode. To enhance the appearance of dim structures, Icy software (66) was used to pseudo-color image channels with the pre defined look-up-table “Fire” based on pixel intensity. Color bars indicate the relative range of pixel intensity (white=high, purple=low, from 0 arbitrary units to 1). Nuclear counterstain (gray) was overlaid as a reference. Images displayed in the ‘Fire’ view, based on a logarithmic scale (“Log Scale”), illustrate both high-intensity and low-intensity granules in the same image.

CHIKV nsP3 sequesters G3BP1/2 in the context of either a replicon (16, 26) or infectious virus (24, 25), thereby interfering with SG responses. Recent sub-diffraction microscopy revealed stable substructures of G3BP1 protein within SGs (29, 30). To determine whether nsP3-granules also sequestered G3BP proteins and contained similar substructures, we imaged stable CHIKV cells with Airyscan microscopy. Airyscan or “Image Scanning Microscopy” (31, 32) relies on array detectors to reassign photon-pixels and oversample the pattern from diffracted light, thereby improving image resolution (by a factor of 1.7) and sensitivity (33). Airyscan outperformed standard confocal microscopy and was sensitive enough to detect small granular structures of nsP3 or G3BP2 (see Fig. S1 in the supplemental material). Whereas nsP3 and G3BP appeared to have a diffuse distribution in confocal images, the improved resolution of the Airyscan microscope uncovered an uneven distribution in a large (1.2 μm diameter) granule, consistent with the presence of substructure (Fig. 2, ROI 1 and Fig. S1, ROI 4). In contrast, small granules (0.2-0.8 μm diameter) lacked substructure (Fig. 2, ROI 2-6). Overall, nsP3 and G3BP2 staining overlapped inside granular structures (Fig. 2). Small clusters of nsP3 (0.2 μm diameter) were ten times less intense than large granules. In summary, sub-diffraction microscopy revealed the co-occurrence of nsP3 and G3BP2 in granules that had a continuous size distribution but were associated with substructures only at larger diameters (i.e., 1.2μm). Small granules (i.e. 0.2 μm) that had a ten-fold lower fluorescence intensity than larger granules (0.4-1.2 μm) were also visible as a result of the increased sensitivity of the Airyscan microscope.

**Figure 2.**
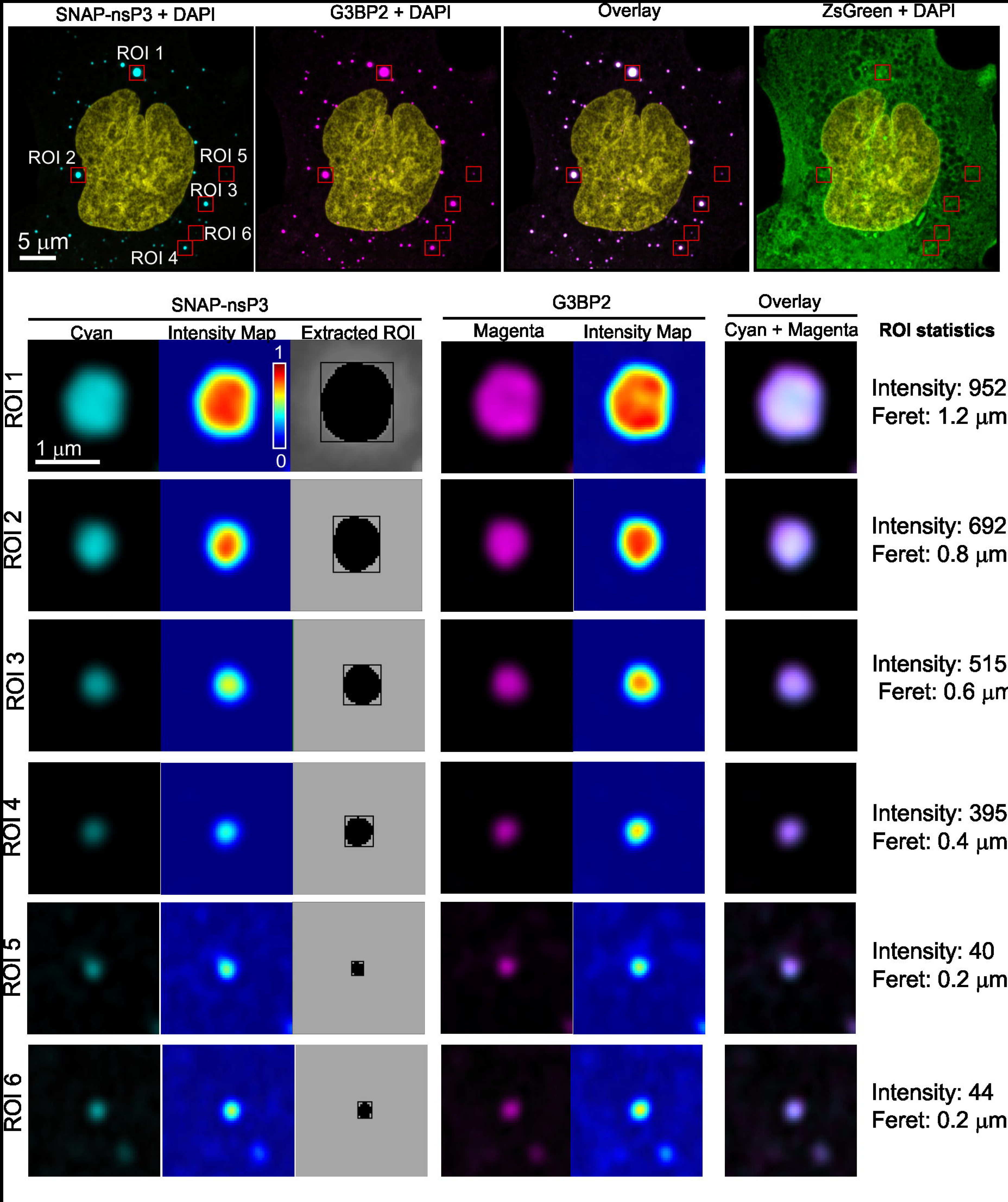
Characterization of nsP3-G3BP interaction by sub-diffraction microscopy. SNAP-nsP3 (cyan) was stained with BG-647 SiR as in Fig. 1 and G3BP2 (magenta) was immunostained with rabbit antibodies that bind to G3BP2 and secondary anti-rabbit IgG antibodies conjugated to Alexa 594. Stable CHIKV cells constitutively expressed ZsGreen (green). Nuclei were counterstained with DAPI (yellow). Images were acquired by Airyscan microscopy operated in super-resolution mode. Top panel shows an overview of the imaged cell with boxed regions-of-interest (ROIs) that are magnified in lower panels (ROI No. 1-6). Overlaps between cyan and magenta layers appear in white (Overlay). Intensity maps were created in Icy software and represent relative pixel intensity according to colormap ‘Jet’. For high-contrast display of nsP3-granules, contrast was optimized within each image by adjusting the view range in the histogram viewer window of Icy software. Outlines of the extracted ROIs that were used for bioimage analysis are shown in grayscale. Intensity: the average intensity distribution in the nsP3 channel inside the ROI (in arbitrary units). Feret: maximum Feret diameter, the maximum distance between any 2 points of the surface. Images represent single slices, which were extracted from Z-stacks.

### Juxtaposition of nsP3-granules, dsRNA foci, nsP1-positive structures, cytoskeleton, early endosomes, and nucleoporin

During the viral life cycle nsP3-granules sequester G3BP, thereby blocking SG assembly (16, 17). The relationship between cytoplasmic nsP3-G3BP complexes and CHIKV RNA synthesis is less clear, as viral dsRNA foci show limited overlap with nsP3-G3BP clusters in cells transfected with a replicon (16) or with nsP1 in infected cells (25). However, more recently it was reported that large cytoplasmic and small plasma-membrane-bound G3BP-nsP3 complexes bind viral genomic RNA during CHIKV infection, with dsRNA-containing viral replication complexes forming nearby (24). The authors suggested that nsP3-G3BP granules play an extra role aside from merely sequestering SG-related proteins.

To further explore the relationship between nsP3 and replication sites in stable CHIKV cells, we used Airyscan microscopy to visualize SNAP-tag labeled nsP3, together with immunostaining for dsRNA and nsP1. Whereas the antibody against dsRNA can identify alphavirus replication complexes (34), the fluorescence of ZsGreen can serve as an indirect readout of the viral subgenomic RNA. Discrete dsRNA-foci were spread throughout the cytoplasm (Fig. S2B). Rather than completely overlapping with larger nsP3-granules, dsRNA foci were in a proximal location and often juxtaposed (Fig. 3). In another example, a dsRNA focus coincided with a smaller nsP3-containing cluster (Fig. 3, Cell 2, ROI 1). Ring-like structures coated with nsP1 were also near these dsRNA foci (Fig. 3). The proximity of dsRNA foci, nsP1-coated structures, and nsP3-granules suggested that the latter not only sequestered G3BP protein but also played a role in viral replication.

**Figure 3.**
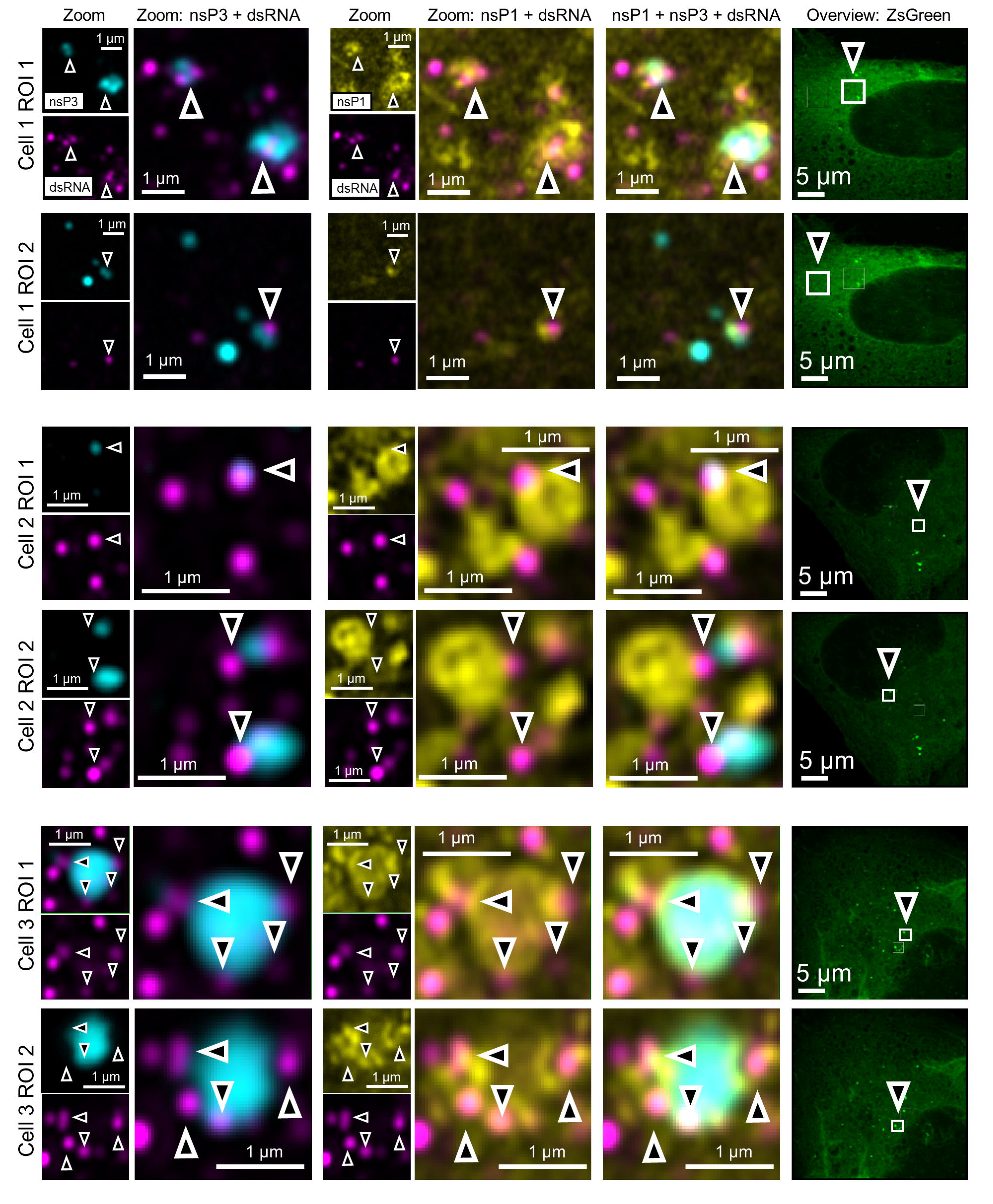
Four-color microscopy of nsP3, dsRNA, nsP1 and ZsGreen. Stable CHIKV cells were fixed and probed for nsP3 (cyan), dsRNA (magenta), nsP1 (nsP1), and ZsGreen (green) by a combination of SNAP-tag labeling and indirect immunofluorescence assays. Images were taken with an Airyscan microscope operated in super-resolution mode. Zoomed-in views taken from Fig. S2B are shown here in panels labeled according to individual cell (Cell 1-3) and ROI (ROI 1-2). Overlay images are a combination of the nsP3, nsP1 and dsRNA layers as indicated. The zoomed-out ZsGreen channel is shown as a separate reference with the corresponding ROI marked by a white box. Arrowheads indicate regions of proximity between nsP3, dsRNA, and nsP1.

As described above, nsP3-containing granules were part of a unique microenvironment that also housed dsRNA foci and nsP1. To further characterize the environment surrounding granules in stable CHIKV cells, we probed for cytoskeletal proteins (vimentin, β-tubulin, β-actin). Stable CHIKV cells had intact networks of vimentin, β-actin, and β-tubulin (Fig. S3, Cells 4-6). Magnifications of nsP3-granules showed they were associated with the cytoskeleton (Fig. 4, arrowheads). Sometimes, patches of vimentin or β-tubulin appeared to partially enclose nsP3-granules (Fig.4, Cell 4 ROI 1-3, Cell 5 ROI 1). Furthermore, a screen with antibody markers of cellular compartments (ER, mitochondria, early endosomes, and the nuclear membrane) showed that nsP3-granules were often closely associated with Rab5 (early endosome marker), and Nup98 (nuclear pore protein) (Fig. 4). NsP3-granules were detected close to Rab5-positive organelles, but were not contained within them, as Rab5 and nsP3 did not overlap (Fig. 4, Cell 7). Granules were also located (i) at the nuclear membrane (Fig. 4, Cell 8 ROI 1 & 2), flanked by Nup98-containing regions; and (ii) near cytoplasmic clusters of Nup98 (Fig. 4, Cell 8 ROI 3-4). In summary, nsP3-containing granules of various sizes interacted with the cytoskeletal network, early endosomes and Nup98-containing structures.

**Figure 4.**
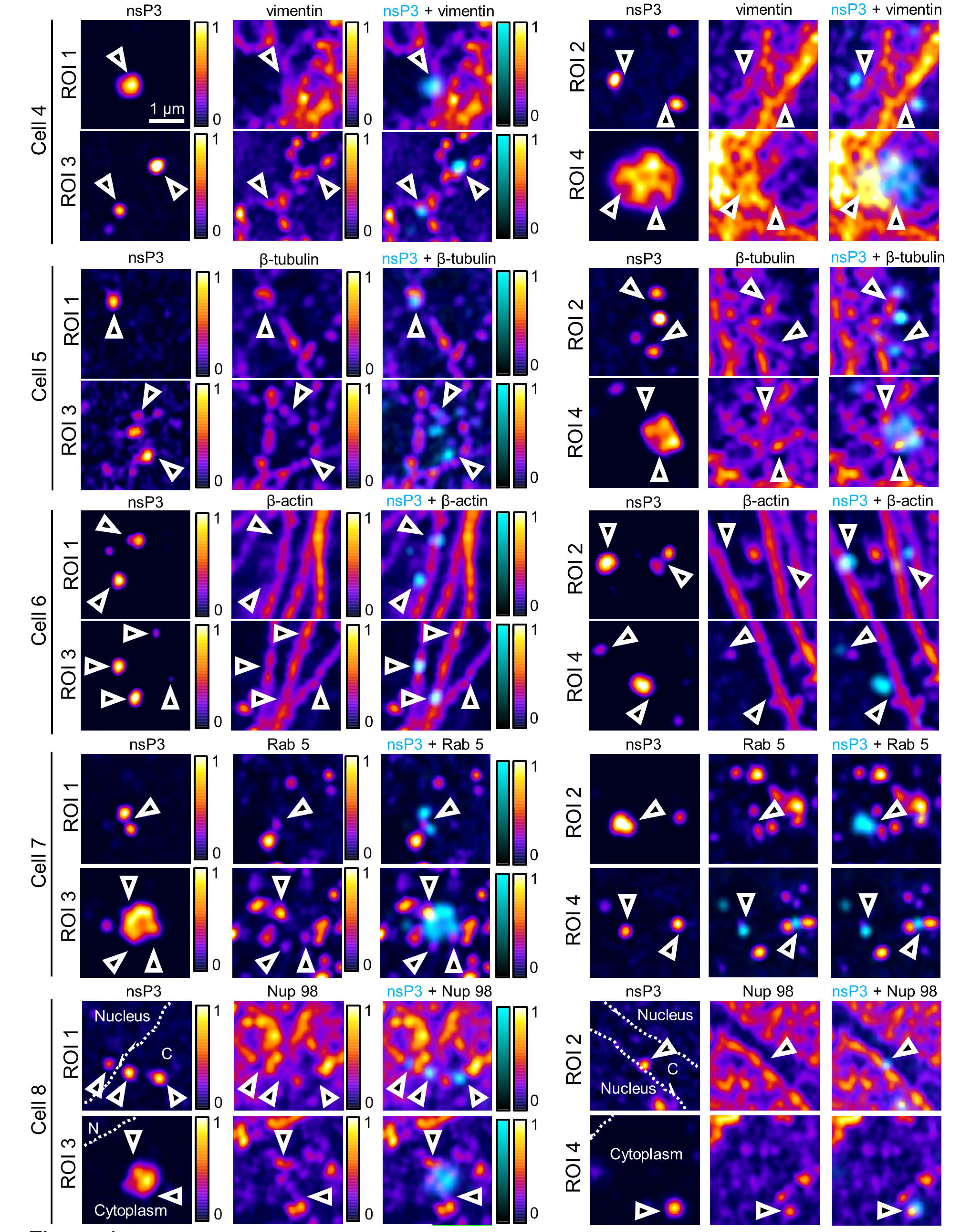
Cellular structures associated with nsP3-granules. Fixed, stable replicon cells were examined for presence of nsP3 and cellular markers by indirect immunofluorescence. The magnified regions-of-interest were derived from imaged cells for which overviews are provided as a reference in the supplemental material alongside corresponding DAPI and ZsGreen layers (Fig. S3). To enhance the appearance of dim structures, single-channel images of nsP3 or Nup98 were both displayed with the “Fire” colormap in Icy software. To distinguish the two channels in overlay images (“nsP3 + “Respective Cellular Marker”), the nsP3-layer was pseudocolored in cyan. Arrowheads serve as digital fiducial markers and point towards regions where nsP3 is associated with a cellular structure. Dashed lines for Cell 8 indicate the nuclear membrane and were drawn according to DAPI images (Fig. S3). Images are single slices extracted from Z-stacks that were taken with an Airyscan microscope operated in super-resolution mode.

### Imaging the dynamics of nsP3-containing granules within stable CHIKV cells

SNAP-reagents can label live cells, allowing both the analysis of movement of tagged proteins, as well as pulse-chase studies to examine protein turnover. Stable CHIKV cells labeled at the onset of a pulse-chase experiment still contained “aged” nsP3-granules after chase periods of 24 h and 48 h (Fig S4A). To track individual granules over time, we also imaged for shorter intervals (every 30 min). Large cytoplasmic nsP3-granules could be monitored over the length of the recording (2 h) and did not visibly disassemble (Fig. S4B). The addition of a non-fluorescent SNAP ligand (=quench) in complementary quench-pulse-chase experiments blocked all binding sites of the SNAP-tagged protein pool (Fig. S4C, FOV1). After a defined chase period of 1 hour in unlabeled media, pulsing with the fluorescent SNAP-reagent uncovered an unblocked population of nsP3-granules, consistent with newly synthesized protein accumulating in granular structures (Fig. S4C, FOV2). The staining of unblocked nsP3-granules increased with a 24-h chase (Fig S4C, FOV3).

To further study the intracellular transport of nsP3-granules, we imaged stable CHIKV cells at 8-s intervals with standard confocal microscopy. This revealed a mixture of nsP3-granules with (i) total displacements <5 μm within perinuclear regions (Table S2, Movie S1 & Fig. 5A, open arrowheads); and (ii) total displacements >7 μm (closed arrowheads). Next, we used instant structured illumination microscopy (iSIM) for live-cell recordings at higher frame rates (35). ISIM increases spatial resolution by a factor of √2 compared with widefield microscopy, and by a further factor of √2 with post-processing, while rapid image capture provides the temporal resolution needed for dynamic events within cells. Whereas small nsP3-granules moved through the cytoplasm over short distances with intermittent bursts of speed (Movie S2 & Fig. 5B, open arrowheads), large granules remained static during the recording and had a low net-displacement (Movie S2 & Fig 5B, closed arrowheads). In summary, the dynamic analysis of nsP3-granules showed that they (i) could persist in cells for days, (ii) accumulated newly synthesized protein and (iii) could be classified into static and motile subclasses with characteristic displacements and speeds.

**Figure 5.**
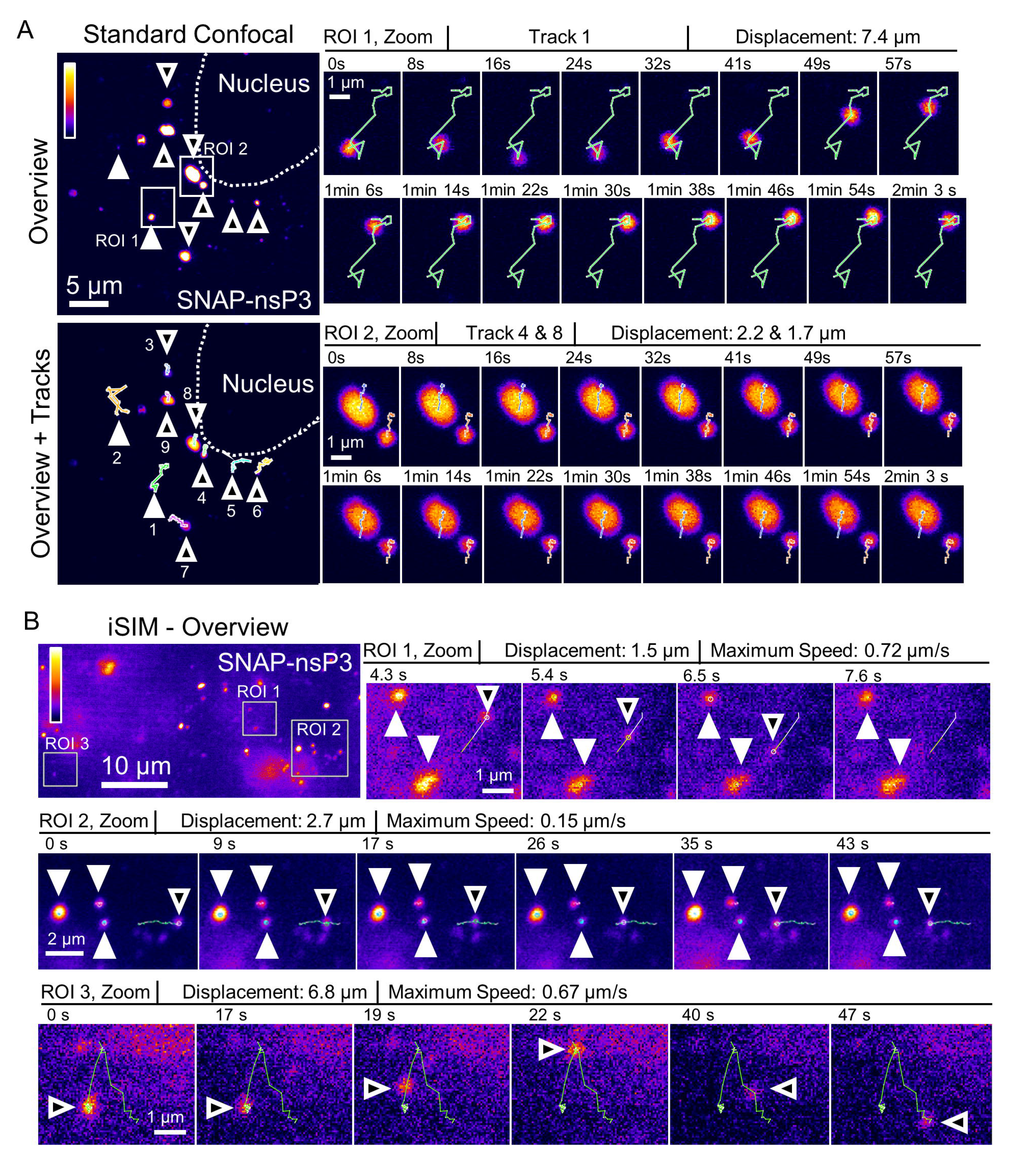
Live imaging of SNAP-nsP3 in stable CHIKV cells showing movement patterns of nsP3-granules. A. Live cells were labeled with BG-647 SiR and examined with an inverted ZEISS LSM 700 confocal laser scanning microscope. A time-lapse series was acquired in the far-red channel (SNAP-nsP3) at intervals of 8.2 s for 20 cycles (=155 s). Overview images on the left represent the first images of the recordings included as Movie S1 in the supplemental material. Positions of individual granules were tracked from frame-to-frame and overlaid on the first image of the recording (“Overview + Tracks”). Numbers 1-9 correspond to track numbers. Closed arrowheads mark granules with higher displacement than that of granules marked by open arrowheads. Single-channel image of SNAP-nsP3 is pseudocolored according to predefined colormap ‘Fire’ in Icy software. A median filter was applied to remove background pixels. The insets depict the paths of SNAP-nsP3 granules in two ROIs. Total displacement of the tracked granule are shown. B. Live imaging of SNAP-nsP3 in stable CHIKV cells by instant structured illumination microscopy (iSIM). Entire recording along with zoomed-in views is also included in Movie S2. The 2-D time-lapse series consisted of 100 frames. Images were acquired at intervals of 1080 msec. Closed arrowheads mark static structures, open arrowheads mark motile structures. Displacement: the sum of all consecutive displacements in each track, which corresponds to the total distance travelled by the granule.

### Static internal architecture of nsP3-granules during persistent replication

To further investigate the dynamics of nsP3-granules we addressed the substructure of individual granules. As a reference, we expressed an EGFP-G3BP1 fusion in Huh-7 cells, selected cells with a diffuse G3BP distribution, treated cells with sodium arsenite to induce SGs, and visualised G3BP-granules by fluorescence recovery after photobleaching (FRAP). EGFP fluorescence recovered within seconds after the photobleach (Fig. 6A, Fig. S5A), consistent with G3BPs rapidly shuttling into and out of SGs. To ask whether nsP3-granules exhibited the same dynamic property, we repeated FRAP experiments in stable CHIKV cells labeled with BG-TMR-Star. No fluorescence recovery or redistribution occurred over the duration of the experiment (Fig. 6B, Fig. S5B), suggesting that nsP3 remained fixed within the granular architecture and did not undergo the dynamic exchange with the surrounding cytoplasm as seen for G3BP.

**Figure 6.**
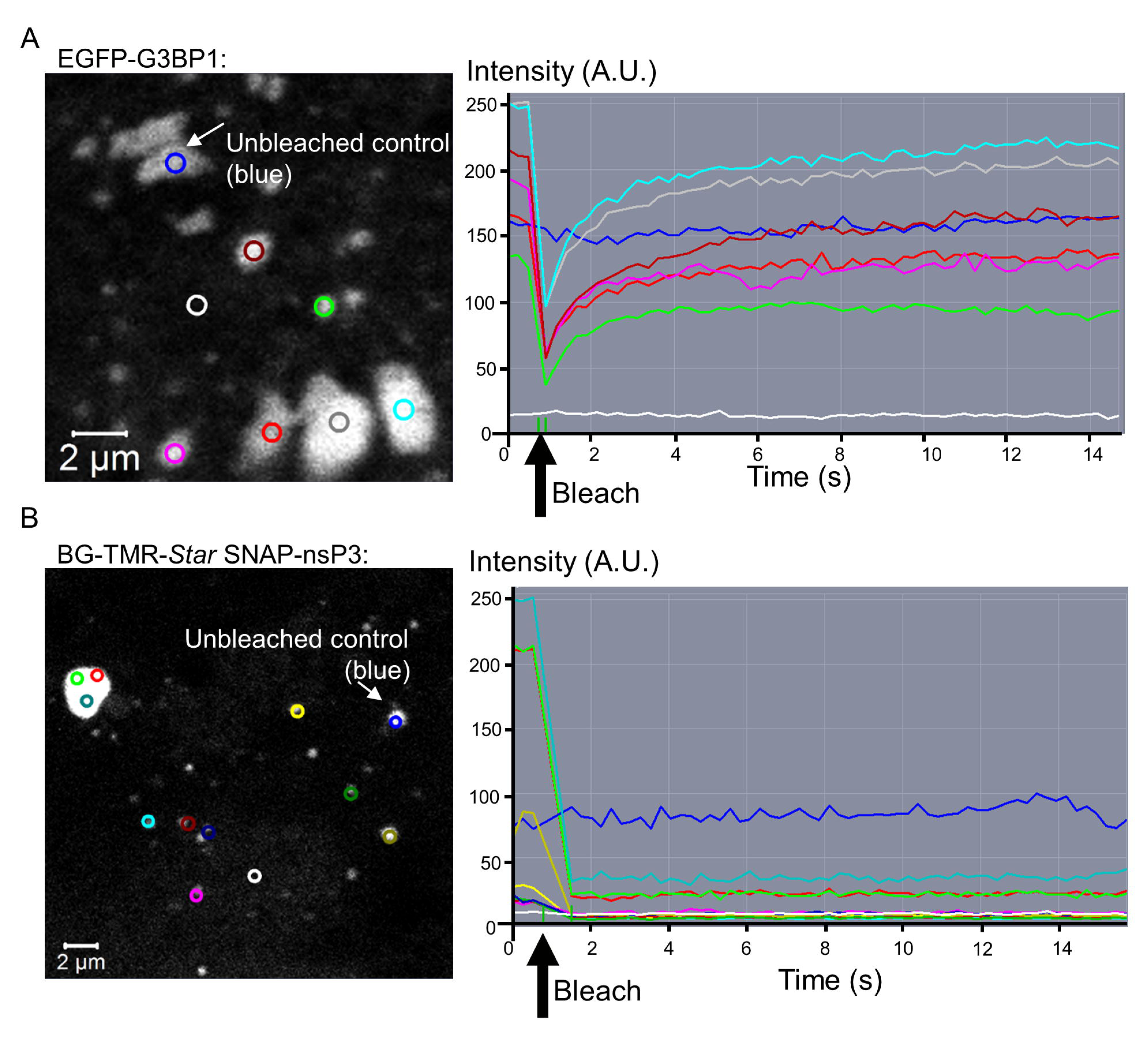
Static internal architecture of nsP3-granules. A. HuH-7 cells were transfected with GFP-G3BP1 plasmid. At 48h post-transfection, cells were examined by live cell imaging with an LSM700 microscope with a Plan-Apochromat 63x NA 1.4 Oil objective. Field-of-views with cells overexpressing GFP-G3BP granules were selected. Scan zoom was set to a factor of 6.6. Images of cells were recorded every 250 ms. Circular regions (circles) were bleached with 488-nm laser pulses after two cycles of imaging. Areas overlapped with part of a large G3BP-granule or an entire granule. A reference area was included that was not photobleached. Grayscale image (left) provides overview of bleached regions (circles). Circle color corresponds to lines in adjacent graphs (right), which plot the mean fluorescence intensity within each bleached area over time. Blue circle marks the unbleached control region. Intensity is measured in arbitrary units (A.U.). B. FRAP of stable CHIKV cells stained with BG-TMR-Star. The same settings shown in A were used, except for bleaching with the 555-nm laser line instead of 488-nm.

Previously, G3BP1-containing SGs were shown to be stable in lysates of stressed cells, suggesting that these membrane-less organelles are made up of stable core structures (29). To test whether this was also the case for nsP3-granules, we lysed stable CHIKV cells and examined the lysates by microscopy. Bright-field images of cell lysates indicated the presence of refractive granules, while fluorescence microscopy identified granules that had incorporated the BG-TMR-Star label (Fig. 7A). Next, we wanted to test whether nsP3 persistence led to an inability to respond to oxidative stress. Stable CHIKV cells maintained high levels of SNAP-nsP3 and ZsGreen for up to two months. In the absence of puromycin selection, however, cells with reduced or undetectable ZsGreen-fluorescence accumulated (discussed in Text S1). These cells were sensitive to puromycin (data not shown), suggesting that they no longer harbored the replicon. As expected, ZsGreen-positive cells sequestered G3BP2 into nsP3-containing granules, and new G3BP2-granules were absent after arsenite-induced stress (Fig. 7B, ROI 1). In contrast, cells lacking ZsGreen were able to form G3BP2-positive clusters after arsenite treatment (Fig. 7B, ROI 2). Therefore, the renewed ability to respond to arsenite-induced stress was associated with a loss of viral replication and nsP3-granules.

**Figure 7.**
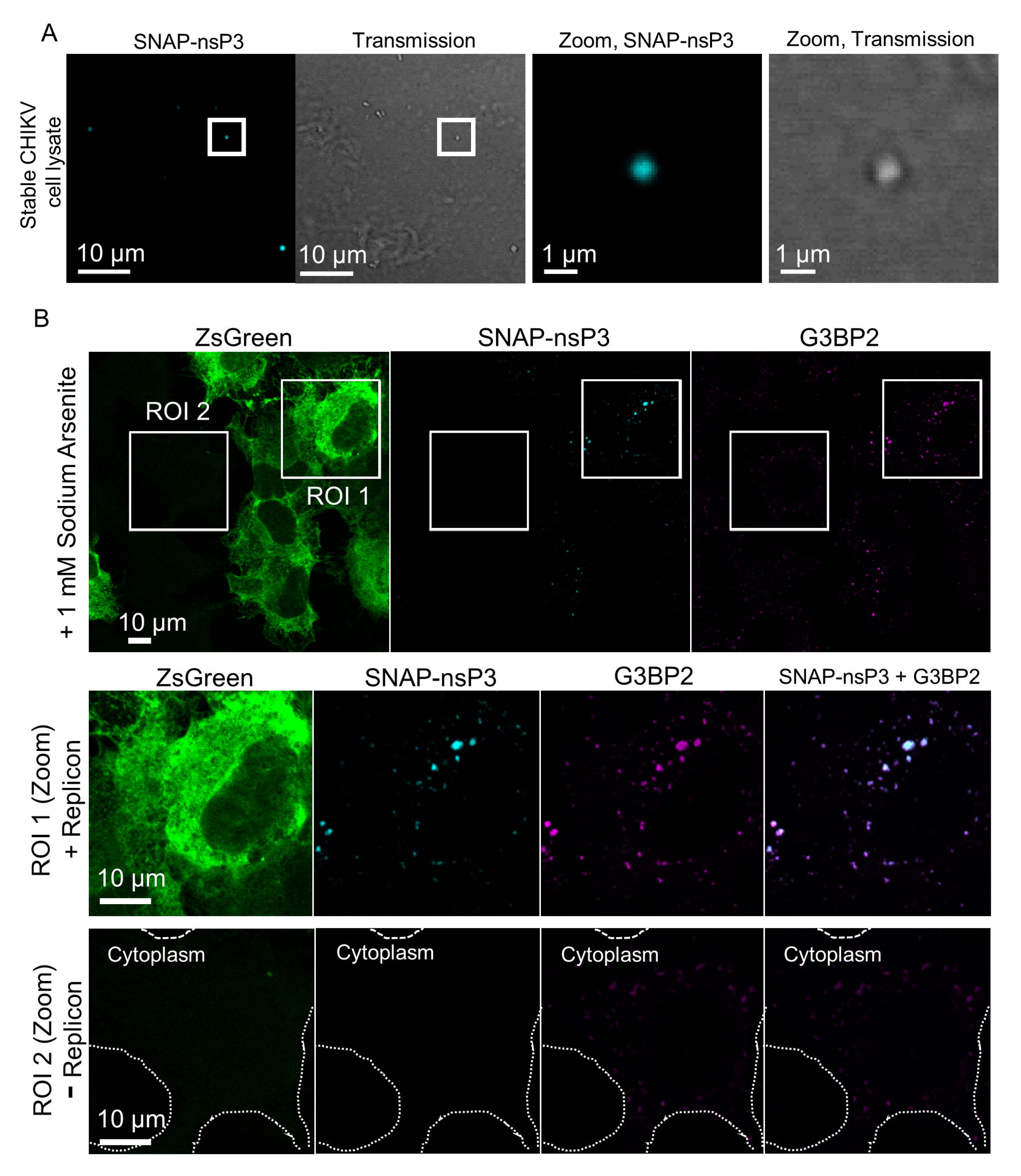
A.Cell lysates from stable CHIKV cells. Live cells were stained with BG-TMR-Star and lysed with Glasgow Lysis buffer. The lysate was then bound to plastic chamber ersistence of NsP3 during Stable CHIKV Replication slides overnight and imaged the following day. Images were acquired with an LSM880 microscopy operated in standard confocal mode. B. Confocal imaging of stable CHIKV cells passaged in the absence of puromycin. Cell population was made up of a mixture of ZsGreen-positive and ZsGreen-negative cells, which only appear in the absence of puromycin. To induce cellular stress granules, sodium arsenite was added for 30 min. Cells were fixed, then stained for SNAP-nsP3 and G3BP2. Stained cells were imaged by standard confocal microscopy. ROI1 is centered on a cell expressing ZsGreen, whereas ROI2 focuses on a cell that is ZsGreen-negative. Cell boundaries appearing as dashed white lines were drawn based on green auto-fluorescence of the cell (only seen when overexposing green channel).

## Discussion

The objectives of this study were firstly to characterize the interaction between CHIKV nsP3 and cellular components during persistent replication, and secondly to evaluate the persistence of cytoplasmic granules composed of viral and cellular proteins. To achieve these objectives, we expanded the utility of a non-cytotoxic replicon by combining it with SNAP-tag-based fluorescent labelling and sub-diffraction multi-color microscopy to provide unprecedented insights into the substructure of persistent nsP3-G3BP-granules. These studies revealed their relationship with dsRNA, nsP1-positive structures, and cellular organelles and examined their dynamics to uncover a stable population of nsP3-granules along with a subclass of nsP3-positive structures trafficking through the cell cytoplasm. Importantly we observed that nsP3-granules lacked a dynamic internal architecture and remained stable in cell lysates. Lastly, we showed that the ability to respond to oxidative stress was associated with the loss of CHIKV replication and nsP3-granules.

### Stable CHIKV cells as a versatile tool for studying cytoplasmic nsP3-granules

Previous reports on non-cytotoxic Old World alphaviruses elucidated the relationship between the loss of cytotoxicity and nsP2-specific mutations that lead to reductions in replication (28, 36–38). The cytopathic effects of a previously described SNAP-tagged replicon limited its study to transient experiments. We now overcome this limitation with a new HuH-7 cell line that harbors replicating CHIKV RNA and encodes both SNAP-tagged nsP3 and ZsGreen as a genetic reporter for viral subgenomic RNA. Whether this replicon only establishes persistent replication in specific cell types as has been observed for other non-cytotoxic replicons (28, 36) remains to be determined. We found that the SNAP-tagged replicon also persisted in C2C12 mouse myoblasts, albeit less efficiently, whereas a glial cell line did not support continuous replication (R. Remenyi, unpublished data).

To our knowledge, the system presented here is the first to allow intracellular tracking of nsP3 during persistent replication of CHIKV RNA. A similar accumulation of nsP3 in cytoplasmic granules occurs in transient replicons (16, 26, 27) and during late stages of infection (24, 25, 27). Strikingly, SNAP-nsP3 in stable CHIKV cells did not form rod-like structures, which were observed in cells infected with CHIKV^SNAP-P3^. An overview of the literature on these rod-like structures and their appearance under different experimental conditions is presented in the supplemental material (Text S1). Surprisingly, infection with CHIKV^P3-ZsGreen^ was not associated with the presence of rod-like structures. However, we cannot rule out that rod-like structures only form transiently and are no longer present at the observed time point.

Nonetheless, the lack of rods was not accompanied by a reduction in infectious titers. Thus, our results suggest that the ability to form rod-like structures can be affected by the sequence of the inserted tag in the C-terminal domain (SNAP vs. ZsGreen), but also whether nsP3 is expressed during persistent replication or infection. Interestingly, the non-cytotoxic replicon also encodes a leucine residue instead of isoleucine at position 175, in a presumed unstructured region between predicted domains of nsP3 (28). Although this mutation may primarily stabilize replication complexes in conjunction with other non-cytotoxic mutations (28), we do not know yet whether it affects the formation of rod-like structures. Taken together, SNAP-nsP3 can form a mixture of cytoplasmic rod-like granular structures during CHIKV infection, but only granules persist in cells that persistently replicate CHIKV RNA.

### Persistence of nsP3-G3BP granules within a microenvironment containing dsRNA, nsP1, and cellular markers

Sub-diffraction multi-color microscopy of stable cells revealed that nsP3-granules were (i) G3BP2-positive (ii) juxtaposed to dsRNA foci and nsP1-positive structures (iii) associated with cytoskeletal markers and (iv) proximal to Rab5-and Nup98-positive organelles. Alphavirus nsP3 forms cytoplasmic granules with vertebrate G3BP and the mosquito homolog Rasputin (16, 17, 21, 24, 25, 39–41). The non-cytotoxic replicon preserved this interaction in cytoplasmic granules whose diameter and protein-content varied. Moreover, only the larger granules (>1 μm) appeared to have an internal substructure. In the future, stochastic optical reconstruction microscopy (STORM), which can provide an even higher resolution to Airyscan microscopy, may be necessary to reveal the detailed substructure of smaller granules (< 500 nm). For example, STORM revealed that G3BP-containing SGs had stable core structures of ~200-nm diameter (29).

Multi-color Airyscan microscopy provided a convenient workflow to examine ZsGreen-expressing stable cells for interactions between nsP3, dsRNA, and nsP1. Alphavirus nsP1 can bind membranes (42, 43) and may use its membrane-binding domain to tether replication complexes to cellular membranes (44). During infection of the related Semliki Forest Virus (SFV), nsP1 co-localizes with G3BPs in putative replication complexes (17). However, the nsP1:nsP3 and nsP1:G3BP association could not be clearly detected during transient CHIKV replication and CHIKV infection (16, 25, 26). We were able to image a partial overlap of nsP1-positive structures with nsP3 granules in stable cells. Occasionally, nsP1 coated ring-like structures, which may represent virus-induced membranous organelles. Furthermore, we could detect dsRNA-positive foci in contact with nsP3-granules. Previous studies outlining the relationship between dsRNA and replication complexes during alphavirus infection are further discussed in the supplemental material (Text S1).

During the viral life cycle, SFV and Sindbis virus (SINV) form cytopathic vacuoles, which measure 0.6-2 μm in diameter. Thus, nsP3-structures that are associated with dsRNA and ring-like structures of nsP1 in this study may be related to cytopathic vacuoles. Ultimately, correlative light and electron microscopy (CLEM) of stable CHIKV cells can elucidate the ultrastructure of nsP3-granules and their relationship with membranous organelles, as was done for Semliki Forest virus (SFV) (45). A useful feature of stable CHIKV cells is the fact that 100% of puromycin-selected cells have ongoing replication and that ZsGreen can serve as a reference for finding the same cell in CLEM approaches.

We also captured high-resolution images of an association between nsP3-granules and cytoskeletal markers. Previous microscopy analysis of vimentin and dsRNA during CHIKV infection implicates vimentin in an anchorage network that supports replication complexes (46). Vimentin also co-localizes with nsP3-containing complexes during SINV infection (22). Likewise, the cellular vimentin scaffold plays a role in directing Dengue virus replication complexes to perinuclear areas via an interaction with the viral NS4A protein (47). Thus, the observed concentration of nsP3-granules in perinuclear regions (Fig. 1, Fig. S1-2) could be explained by an nsP3:vimentin interaction, previously identified in proteomic studies (46). We note that a further discussion of known interactions of alphaviruses with actin and tubulin networks, as well as Rab5-endosomes is provided in the supplemental material (Text S1). Our data provide evidence for similar interactions during persistent replication.

We also investigated the previously unexplored relationship between nsP3 and the nucleoporin Nup98. Little is known about the nuclear transport of nsP3, while the localization of nsP2 to the nucleus is well-documented (16, 40, 48, 49). Intriguingly, a role for G3BP as a nuclear transport factor has been proposed and SINV nsP3 has been identified at the nuclear membrane (21). Our results imply that nsP3-granules associate with a nucleoporin during persistent replication and may connect to RNA transport pathways at the nuclear membrane. Viral proteins that bind to Nups or RNA transport factors have been shown to stimulate remodeling of the nuclear membrane and affect nuclear transport of cellular mRNA and proteins (50, 51). During SFV infection, many nuclear proteins re-locate to the cytoplasm where they play both pro-viral and anti-viral roles (52). We also observed an association of nsP3-granules with cytoplasmic Nup98. During HCV infection, cytoplasmic nucleoporins accumulate at sites rich in viral proteins, including virus-induced membranous organelles and cytosolic lipid droplets (53, 54). In summary, Nups may play a role in persistent replication of CHIKV, which could hijack the physiological functions of nucleoporins to transport CHIKV nonstructural protein components, mRNA, viral RNA, or cellular proteins. Our data warrant a further investigation of this hypothesis.

### Stable CHIKV cells contain mixture of static and dynamic nsP3-granules, which lack dynamic internal architecture and are stable in cell lysates

A key feature of the SNAP-tag is the experimental control over the time-of-labeling, enabling studies protein turnover. NsP3-granules were stable for hours and persisted for days. Granules were also the site where newly synthesized nsP3 accumulated. Thus, old and new populations of nsP3 may continuously mix within cytoplasmic granules, as was seen during transient replication (26). Live-cell microscopy also provided the first real-time tracking of CHIKV nsP3-granules and in-depth view of granule dynamics in mammalian cells. An overview of previous live-cell microscopy results, which identified subclasses of nsP3-structures based on movement patterns in SFV-infected cells (55), is provided in the supplemental material (Text S1). We report similar movement patterns, including (i) the presence of immobile granules within perinuclear regions and (ii) granules moving over short (1-2 μm) and long distances (>7 μm) at maximum speeds between 0.2 and 0.7 μm/s. Because some of these patterns suggest actin-based transport, we are currently setting up additional live-cell microscopy experiments with live-cell probes for actin, but also microtubules or secretory vesicles. In the future, stable CHIKV cells can provide invaluable real-time insight into interactions between CHIKV and the host through multi-color imaging of ZsGreen, far-red-fluorescent SNAP-nsP3 labeling, and a third, blue- or red-fluorescent marker.

FRAP experiments revealed the static internal architecture of nsP3-granules, whereas arsenite-induced G3BP-granules had a similar fluorescence recovery as seen in human osteosarcoma cells (56). The absence of a rapid exchange in CHIKV-induced granules implies that nsP3 may play a role that differs biochemically from the dynamic role of G3BP in SGs (29, 30, 57). For example, nsP3 may create a scaffold similar to the one formed by Fas-activated serine/threonine kinase (FASTK) in SGs (57). Although we cannot rule out that G3BP shuttles in and out of nsP3-granules, we predict that G3BP would be fixed in granules in a similar way: nsP3 completely overlapped with G3BP and nsP3-granules were stable enough to be preserved in cell lysates. Moreover, previous studies demonstrated that alphavirus nsP3-G3BP granules lack canonical SG markers (16, 17) and remain stable during cycloheximide treatment (16), which dissolves SGs (58). FRAP experiments of membrane-associated foci containing non-structural proteins of another RNA virus, HCV, also found a limited exchange between clusters of non-structural proteins and the periphery (59–61). Thus, some of the nsP3-structures may represent cytopathic vacuoles, in which nsP3 has a limited exchange with the surrounding cytoplasm.

Unlike cytopathic vacuoles, which would be sensitive to detergents, a population of nsP3-granules was detergent-resistant and stable in cell lysates. This persistence in lysates mimics that of mammalian SG cores (29). Jain et al. describe a dynamic shell around core structures that gives SGs biochemical qualities akin to liquid-liquid phase separations. We propose that similar stable core structures could make up nsP3-G3BP granules. Text S1 provides a brief overview of the link between liquid-liquid phase separation, membrane-less organelles, stress responses, and toxic protein clusters, which are a hallmark of neurodegenerative disease. This link forms the basis of a new hypothesis that nsP3-granules can perturb cellular responses to environmental conditions. However, more experiments are needed to (i) further characterize persistent nsP3-granules biochemically (ii) identify other cellular or viral proteins within granules; and (iii) induce granular disassembly. Clearing cells of these stable cytoplasmic complexes could be essential for preventing toxic effects that only emerge during prolonged exposure to CHIKV proteins. Moreover, directly targeting persistent nsP3-granules could lead to new approaches to combat Chikungunya virus infections. Encouragingly, cells that cleared the CHIKV replicon during culturing in the absence of selective pressure were able to regain the ability to form SGs in response to arsenite treatment.

In summary, our results present the first evidence that granules containing the viral protein nsP3 and cellular protein G3BP persist in human cells with autonomously replicating CHIKV RNA. Generation of a cell line harboring a persistently replicating SNAP-tagged replicon and advances in microscopy technology allowed us to reveal interactions between SNAP-nsP3, viral components (nsP1, dsRNA), and cellular components (cytoskeleton, endosomes, nucleoporin). Overall, nsP3-granules were stable, differed in their mobility, lacked a dynamic internal architecture, and were stable in cell lysates. These findings may also have clinical relevance, as CHIKV can cause chronic infection and persist in various cell types, such as macrophages, muscle, and liver cells. However, whether prolonged exposure to nsP3-granules causes pathogenic changes within the cell and can contribute to the long-term effects of Chikungunya disease remains to be determined in future studies. Lastly, the reagent presented in this study adds a new dimension for future explorations of host-pathogen interactions, in particular as they relate to nsP3, and for the search for inhibitors that specifically target nsP3.

## Materials and Methods

### CHIKV constructs

The replicon CHIKVRepRLuc-FL-5A-PG-IL was described previously and allows for stable, non-cytotoxic growth in HuH-7 cells (28). It contained a cassette encoding a puromycin-N-acetyltransferase (Pac) - FMDV 2A autoprotease - ZsGreen fusion under the control of the sg-promoter. Further information on the construction of a SNAP-tagged derivative of this non-cytotoxic replicon and infectious clones CHIKV^SNAP-nsP3^ and CHIKV^ZsGreen-nsP3^ is provided in the supplemental material (Text S2).

### Cells, media, transfection, and infection

HuH-7 cells were maintained in complete media (Dulbecco's modified Eagle's medium supplemented with fetal calf serum, penicillin, streptomycin, non-essential amino acids and HEPES Buffer) as described previously (26). HuH-7 is a well differentiated hepatocyte-derived cellular carcinoma cell line taken from the liver tumor of a male Japanese patient in 1982 (62); these cells were from John McLauchlan (Centre for Virus Research, Glasgow). Growth media supplemented with puromycin (final concentration, 5 μg/ml) was used for antibiotic section.

Additional information on electroporating HuH-7 cells with in vitro transcribed RNA of the SNAP-tagged non-cytotoxic replicon can be found in the supplemental material (Text S2). Electroporated cells were seeded in 10-cm dishes. Cells were incubated in puromycin-free media for a minimum of two days before starting puromycin selection. During puromycin selection, cells were monitored with a widefield fluorescence microscope and a FITC filter setup for ZsGreen fluorescence. After ZsGreen-positive cells reached a high proportion (2-5 days), cells were expanded in puromycin-free media. Heterogeneous populations of ZsGreen-positive cells, which we call “stable CHIKV cells”, were collected from confluent T75 flasks to make frozen cell stocks in fetal calf serum supplemented with 10% DMSO (about two weeks after electroporation). At the same time, stable CHIKV cells were passaged under routine conditions and used in microscopy experiments.

### Indirect immunofluorescence assay, intracellular SNAP-tag labeling, and pulse-chase experiments

Primary and secondary antibodies used in indirect immunofluorescence assays (IFA) are listed in Text S2.For intracellular SNAP-tag labeling, stable CHIKV cells were plated in 24-well plates containing 13-mm glass coverslips. The next day, SNAP-nsP3 was fluorescently labeled in live cells as described previously (26, 63) and outlined in Text S2. Note that for staining of SNAP-nsP3 from a viral infection (Fig. 1), infected cells were first fixed at room temperature with 4% formaldehyde for 30 min to inactivate infectious virus, followed by the same SNAP-staining protocol outlined below, since chemical fixation did not abolish the ability of SNAP ligands to bind to the SNAP sequence.

For IFA and staining with G3BP2, β-actin, or J2 antibodies, formaldehyde-fixed cells were permeabilized with 100% Methanol for 10 min at −20°C. For all other antibodies, cells were permeabilized with a buffer containing 5% fetal calf serum and 0.3% Triton × 100. Cells were incubated with primary antibody solution containing 1% bovine serum albumin (BSA) overnight at 4°C, except the mouse J2 antibody, which was incubated for 2 h at room temperature in diethylpyrocarbonate-treated PBS. After three washes in PBS, secondary antibody (anti-rabbit Alexa Fluor 594-conjugated IgG or anti-mouse Alexa Fluor 594-conjugated IgG, Molecular Probes) was added. For nsP3/J2/nsP1 triple staining, rabbit nsP1 antibody was added overnight at 4°C to cells already stained with BG-647 SiR (benzylguanine-silicon-rhodamine) and mouse J2. The following day, cells were washed three times in PBS and secondary antibody (anti-rabbit Alexa Fluor DyLight 405) was added. These cells were not counterstained with 4′,6-diamidino-2-phenylindole (DAPI). However, where indicated (Fig. 1, 2, 4), DAPI was added to visualize nuclei. Coverslips were mounted onto glass slides by addition of ProLong Diamond Antifade Mountant (Molecular Probes).

Protocols for pulse-chase experiments and analyses of protein turnover with fluorescence microscopy are available in the supplemental material (Text S2).

### Sub-diffraction light microscopy

An LSM880 upright confocal microscope with Airyscan (ZEISS) was used to acquire sub-diffraction microscopy images as described previously (26, 63). This microscope provides a maximum lateral resolution of 140 nm and axial resolution of 400 nm for a fluorophore emitting at 480 nm. Microscope settings during the acquisition of a series of axial images (Z-stack) are provided in the supplemental material (Text S2). To increase signal-to-noise ratio and resolution, image stacks were processed by Airyscan batch processing within Zen Black. Single-slice images were extracted to produce panels in Fig. 2, 3 and 4. Cell overviews are provided as maximum-intensity projections in Fig. S2 and 3. Note that although live-cell Airyscan microscopy is technically possible, we only used Airyscan microscopy for fixed cells, since our microscope was not set up for live-cell imaging.

### Live-cell microscopy of stable CHIKV cells

Live-cell imaging was done on an LSM700 AxioObserver inverted confocal microscope (ZEISS) equipped with Plan-Apochromat 63x/1.4 Oil Ph3 M27 objective and an incubator box and heated stage set to 37°C. Cells were grown in 35 mm glass (No. 1.5) bottom dishes with 27 mm viewing area (Nunc). Stable CHIKV cells were stained with BG-SiR and Mitotracker Red FM (Molecular Probes), then maintained at 37°C in an optically clear, physiological, and CO_2_-independent imaging buffer (Molecular Probes, Live Cell Imaging Solution supplemented with 10% fetal calf serum, non-essential amino acids and buffered with 10 mM HEPES). To suppress photobleaching, ProLong Live Antifade Reagent was added according to manufacturer’s instructions (Molecular Probes). The setup of microscope settings is noted in the supplemental material (Text S2).

A home-built instant structured illumination microscope (iSIM) was used to acquire additional time-lapse series (Fig. 5B). This instrument is fitted with an Olympus Water Immersion Objective 1.2 NA UPLSAPO 60XW, and 488Ԕnm and 561 Ԕnm lasers (64). Stable CHIKV cells were stained with red-fluorescent BG-TMR-Star before image acquisition. The heated stage was set to 37°C. The same live cell imaging media described above was used, supplemented with ProLong Live AntiFade reagent. Regions of interest were found using the live iSIM display in the green channel (ZsGreen) to avoid bleaching of the red channel (nsP3). Time-lapse series were acquired by taking images of the red and green channels at intervals of 1080 msec. Note that green and red channels are shown in Movie S2, whereas Fig. 5B only shows the nsP3-channel to emphasize the movement of nsP3-granules (red channel).

### FRAP analysis

To induce genuine stress granules in HuH-7 cells, the plasmid pEGFP-G3BP (kindly provided by Richard Lloyd, Baylor), encoding a EGFP-G3BP1 fusion protein (65), was transfected with Lipofectamine 2000 reagent (Thermo Fischer Scientific) into cells plated in a 35 mm glass (No. 1.5) bottom dishes with 27 mm viewing area (Nunc). After 24 hours, cells containing G3BP granules were identified by live-cell microscopy on a LSM700 confocal system set to 37°C. Stable CHIKV cells, stained with BG-TMR-Star with the live-cell protocol, were used for experiments imaging SNAP-nsP3. Experimental details for fluorescence recovery after photobleaching (FRAP) experiments are provided in the supplemental material (Text S2).

### Isolation of SNAP-nsP3 from cell lysates

Stable CHIKV cells grown in 6-well plates were labeled with BG-TMR-Star according to the live-cell staining protocol outlined above. Cells were collected by scraping into PBS using plastic cell scrapers, followed by centrifugation in 1.5-ml microcentrifuge tubes. Cell pellets were lysed with 300 μl ice-cold Glasgow Lysis Buffer [1% Triton X-100, 120 mM KCl, 30 mM NaCl, 5 mM MgCl_2_, 10% glycerol, and 10 mM piperazine-*N,N*′-bis(2-ethanesulfonic acid) (PIPES)-NaOH, pH 7.2] containing protease inhibitors. Lysates were vortexed for 30 s for four cycles and returned to ice in between cycles. A final spin at 850 g was included to remove remaining cellular debris. The final supernatant was added to a 2-well Ibidi plastic slide with Ibitreat surface for optimal cell adhesion (Ibidi). After an overnight incubation at 4°C, 1 ml of 4% Formaldehyde was added to each well for 1 h at room temperature. Wells were washed with PBS and images with an LSM880 system operated in confocal mode.

### Bioimage analysis

Microscopy images were processed on the Icy (http://icy.bioimageanalysis.org) platform (66). Contrast was optimized in individual images by dragging the adjustable bounds of the histogram viewer, which enhances the contrast in each channel without altering the data (66). Color maps (Cyan, Magenta, Green, Gray, Yellow, Fire or Jet) were applied with the lookup table manager to each channel in combination with the corresponding histogram bounds. Further analyses with the Icy software, including segmentation and tracking of granules are described in Text S2.

## Funding Information

This work was funded by a Wellcome Trust Investigator Award to MH (WT 096670). Purchase of shared equipment was made possible by a Wellcome Trust Multi-user equipment award (Zeiss LSM880 instrument, WT104918MA,“Multifunctional imaging of living cells for biomedical sciences”) and a Royal Society equipment grant to Dr. Jamel Mankouri (Zeiss LSM700 instrument, grant number RG110306). Work to build the iSIM system was supported by MRC grant ref MR/K015613/1. YG was supported by a China Scholarship Council/University of Leeds PhD studentship.

## Acknowledgements

We thank Dr. Sally Boxall and the Bio-imaging Facility within the Faculty of Biological Sciences of the University of Leeds for access and help with Airyscan microscopes and Dr. Jamel Mankouri for access and help with the LSM700 confocal system. We also thank Grace C. Roberts and Raymond Li for assistance in culturing of the HuH-7 cell line and stable CHIKV cells.

## Other information

Competing financial interests: The authors have declared that no competing interests exist.

## Supplemental Material

**Text S1.** Supplemental text.

**Text S2.** Supplemental methods and materials.

**Table S1.** Viral titers of CHIKV^SNAP-P3^ and CHIKV^ZsGreen-P3^.

**Table S2.** Quantitation of tracks.

**Fig. S1. Comparison of confocal with Airyscan images.** Airyscan provided improved resolution and signal-to-noise ratios (SNRs), allowing us to spot clusters of nsP3 inside stable CHIKV cells with unprecedented sensitivity. This is most apparent in magnifications of nsP3-and G3BP2-clusters. Airyscan was able to image faint clusters that did not resolve well with standard confocal microscopy. Whereas some clusters appear to be just one object in the confocal images, Airyscan images confirmed the presence of two clusters close together (ROI 1, nsP3). In this paper, we use the term “granules” for these protein clusters. Airyscan also revealed sub-structures within large granules that were not apparent with confocal imaging. Samples were prepared as described for Fig. 2.

**Fig. S2 Subcellular localization of SNAP-nsP3 during CHIKV^SNAP-P3^ infection and overview of fluorescence staining (SNAP-nsP3, dsRNA, nsP3) in stable CHIKV cells.** (A) Samples were prepared and imaged as described in Fig. 1 by Airyscan microscopy. Maximum-intensity-projections of Z-stacks acquired with Airyscan microscopy. NsP3-channel was pseudocolored with the look-up-table "Fire” in Icy software. Nuclear counterstain (gray) was overlaid as reference. Images displayed in the ‘Fire’ view, based on a logarithmic scale (“Log Scale”), illustrate both high-intensity and low-intensity structures in the same image. (B) Reference overviews of stable CHIKV cells that were imaged with four-color Airyscan microscopy. Samples were prepared and imaged as described for Fig. 3. SNAP-nsP3 signal was pseudocolored with a cyan look-up table and displayed on a logarithmic scale (“Log Scale”) to illustrate both high-intensity and low-intensity structures in the same image. Boxed regions were magnified in Fig. 3.

**Fig. S3 Overview of fluorescence staining (SNAP-nsP3 and cytoskeleton, Rab5, Nup98) of stable CHIKV cells.** Reference overviews of stable CHIKV cells that were imaged with four-color Airyscan microscopy. Samples were prepared and imaged as described for Fig. 4. SNAP-nsP3 signal was pseudocolored with a cyan look-up table and displayed on a logarithmic scale (“Log Scale”) to illustrate both high-intensity and low-intensity structures in the same image. Boxed regions were magnified in Fig. 3.

**Fig. S4 A.Long-term imaging of SNAP-nsP3, live-cell confocal microscopy, and quench-pulse-chase of stable CHIKV cells.** (A) Stable CHIKV cells were plated in glass bottom dishes. Labeling with BG-SiR was carried out the following day. Cells were imaged with a Delta Vision Widewield Deconvolution Microscope with the incubator box set to 37°C. After images were acquired for respective time point, cells were returned to regular cell-culture incubators. After 24 and 48 hours, the same grid positions were found using transmission images and fluorescent signals were imaged with the same settings as the 0-h time point. (B) Live-cell confocal microscopy of stable CHIKV cells. Stable CHIKV cells were plated in glass bottom dishes. The following day, SNAP-nsP3 was labeled with BG-SiR and mitochondria were stained with MitoTracker Red FM. Cells were imaged on an LSM700 AxioObserver inverted confocal microscope (ZEISS) equipped with Plan-Apochromat 63x/1.4 Oil Ph3 M27 objective and an incubator box and heated stage set to 37°C; Z-stacks (11-13 Slices) were acquired every 30 min for a total duration of 2 hours. ZsGreen was pseudo-colored in yellow to contrast the cyan overlay of nsP3 and magenta overlay of mitochondria. (C) Quench-pulse-chase experiment. Stable CHIKV cells were plated in 24-well plates containing glass coverslips. The next day, non-fluorescent bromothenylpteridine was used to block the reactivity of intracellular SNAP-nsP3. Blocked cells were fixed with 4% formaldehyde at indicated times post-block (0 h, 1 h, 24 h) and newly synthesized SNAP-nsP3 was stained with BG-SiR post-fixation. Stained samples were imaged with a LSM880 system (ZEISS) operated in confocal mode. One representative field-of-view (FOV) is shown from each sample. The same laser power and detector settings were used to image each FOV. Z-stacks were acquired to capture all the granules present within cells. Images are maximum-intensity projections. SNAP-nsP3 channel was pseudocolored with the ‘Fire’ look-up table.

**Fig. S5. FRAP of EGFP-G3BP1 and SNAP-nsP3.** Individual images of FRAP experiment shown in Fig. 5 (A) Single-channel images or G3BP and (B) of SNAP-nsP3 were pseudocolored according to predefined colormap ‘Fire’ in Icy software. Boxed regions were magnified within insets (‘Zoom’).

**Movie S1. Time-lapse of SNAP-nsP3 in stable CHIKV cells by standard confocal microscopy and instant structured illumination microscope.** Time-lapse movie of ZsGreen (green), MitoTracker Red FM (magenta), and BG-SiR staining of SNAP-nsP3 (fire) in stable CHIKV cells. Individual frames from these live-cell recordings are shown in Fig. 5A. This is followed by a time-lapse movie of ZsGreen (green) and BG-TMR-Star staining of SNAP-nsP3 (fire) in stable CHIKV cells by iSIM. Images were taken every 1080 msec for 100 cycles, alternating in the green and red channel. Individual frames from these live-cell recordings are shown in Fig. 5B.

